# Cholinergic modulation of interhemispheric inhibition in the mouse motor cortex

**DOI:** 10.1101/2024.02.05.579044

**Authors:** Takashi Handa, Qing Zhang, Hidenori Aizawa

## Abstract

Interhemispheric inhibition (IHI) of the homotopic motor cortex is believed to be effective for accurate unilateral motor function. However, the cellular mechanisms underlying IHI during unilateral motor behavior remain unclear. Furthermore, the impact of the neuromodulator acetylcholine (ACh) on IHI and the associated cellular mechanisms are not well understood. To address this knowledge gap, we conducted recordings of neuronal activity from the bilateral motor cortex of mice during the paw-reaching task. Subsequently, we analyzed interhemispheric spike correlation at the cell-pair level, classifying putative cell types to explore the underlying cellular circuitry mechanisms of IHI. We found a cell-type pair-specific enhancement of the interhemispheric spike correlation when the mice were engaged in the reaching task. We also found that the interhemispheric spike correlation was modulated by pharmacological ACh manipulation. The local field responses to contralateral excitation differed along the cortical depths, and muscarinic receptor antagonism enhanced the inhibitory component of the field response in deep layers. The muscarinic subtype M2 receptor is predominantly expressed in deep cortical neurons, including GABAergic interneurons. These results suggest that GABAergic interneurons expressing muscarinic receptors in deep layers mediate the neuromodulation of IHI in the homotopic motor cortex.

## Introduction

Early physiological studies demonstrated interhemispheric inhibition (IHI) in the homotopic motor cortex, which is the inhibition or attenuation of activation in the motor cortex by prior unilateral activation in the contralateral motor cortex, leading to the attenuation of the output signal (Ferbert et al. 1992; Netz et al. 1995). It is widely believed that the role of IHI in unilateral motor execution is to suppress the neuronal activity of the ipsilateral motor cortex to enable accurate unilateral motor control (Chiarello and Maxfield 1996). The neural circuitry mechanisms underlying IHI involve the corpus callosum, which predominantly consists of transcallosal axons of pyramidal neurons, allowing the interaction of the homotopic cortical regions between hemispheres and GABAergic interneurons in the contralateral homotopic area that receive transcallosal excitatory inputs (Toyama et al. 1974; Kawaguchi 1992; Conti and Manzoni 1994; Palmer et al. 2012; Anastasiades et al. 2018). Otherwise, involuntary mirrored bilateral movements may emerge, as reported in patients with agenesis of the corpus callosum (Marsh et al. 2018). Despite their functional importance in motor execution, the cellular mechanisms underlying unilateral motor function remain unclear.

Cholinergic neurons in the basal forebrain innervate the neocortex. Cholinergic neurotransmission acts diffusely in a wide variety of brain regions, including the motor cortex, as a neuromodulatory system (Kitt et al. 1994; Woolf and Butcher 2011; Do et al. 2016). Cholinergic modulation is associated with the regulation of arousal and plasticity (Gu 2002; Sarter et al. 2003; Woolf and Butcher 2011). Indeed, the cholinergic system is essential for skilled motor learning, which is associated with the reorganization of the motor cortex (Conner et al. 2003; Ramanathan et al. 2009; Ren et al. 2022) and functional recovery from motor deficits after brain injury (Conner et al. 2005). Recent studies have suggested that motor-related axonal activity of basal forebrain cholinergic neurons in the motor cortex (Ren et al. 2022) and the spatiotemporal patterns of acetylcholine (ACh) release correlate with different motor behaviors (Lohani et al. 2022). Considering the known effect of ACh on the modulation of motor behaviors as reported in these previous publications, it is tempting to consider the possible role of ACh in the modulation of IHI.

In this study, we address the role of cholinergic modulation in the cellular circuitry mechanism underlying IHI. To this end, we investigated the characteristics of neural activity in the bilateral motor cortex during a reaching task and their correlation between hemispheres in freely moving mice using electrophysiological recordings. The effect of cholinergic manipulation on the neural correlation between the hemispheres and interhemispheric interactions was examined using electrophysiological recordings combined with pharmacological and/or optogenetic approaches. To further investigate the cellular elements underlying cholinergic modulation of IHI, we analyzed cell subtype-based mRNA expression in the mouse motor cortex and cortical depth-dependent expression of muscarinic cholinergic receptors using immunohistochemical methods. By classifying putative cell types in both the ipsilateral and contralateral motor cortical areas, we found that reaching-related spike activity was altered by paw laterality and reaching task-dependent interhemispheric spike correlation. The interhemispheric spike correlation was oppositely affected by the pharmacological blockage of ACh binding to muscarinic receptors and the breakdown of ACh in the brain. The local pharmacological blockage of ACh binding to muscarinic receptors enhanced IHI in the deep layers of the motor cortex, and muscarinic receptor expression in GABAergic interneurons was observed in the deep layers. These results suggested that ACh could function as a disinhibition mechanism through IHI in deep-layer muscarinic receptor-positive interneurons.

## Materials and Methods

### Animal

All experimental procedures were approved by the Institutional Experimental Animal Committee of Hiroshima University (approval number: A22-66). To prepare the Emx1-Cre;ChR2 mouse line, the Emx1-Cre knock-in mouse line (Kassai et al. 2008) was crossed with Ai32 B6.Cg- *Gt(ROSA)26Sor^tm32(CAG-COP4*H134R/EYFP)Hze^*/J (JAX:024109, The Jackson Laboratory, U.S.A.). To investigate the neural response to interhemispheric input, Emx1-Cre;ChR2 mice were used for electrophysiological recordings following optogenetic stimulation of the contralateral hemisphere (N = 6 for cholinergic receptor pharmacological experiments). C57BL/6J mice (Japan CLEA, Japan) were used for behavioral training, followed by chronic electrophysiological recordings (N = 6, 8–12 weeks old, males), acute electrophysiological experiments with systemic drug administration (N = 6, 16–20 weeks old, males), and immunohistochemistry (N = 3, 12–20 weeks old, females). The mice were housed in group cages with free access to food and water and a 12-h light/dark cycle at controlled temperature and humidity.

### Fabrication of microdrive driving tetrode array

Computer-assisted design of the microdrive were made in Fusion 360 (Autodesk Inc., U.S.A.) based on a previous publication (Voigts et al. 2013) (flexDrive, OpenEphys, https://open-ephys.org/flexdrive). The electrodes were inserted by −250 µm per single turn of the screw embedded in the microdrive printed using stereolithography three-dimensional (3D) printer (ProJet HD3000, 3D systems Inc., U.S.A.). We designed the main body (driving electrodes) and base plate (specifying the target in the brain) independently to enable the targeting of a variety of combinations of brain regions. We fabricated tetrode which consists of four NiCr wires (12 µm in diameter, Sandvik AB, Stockholm) with an apparatus to twist the wires (Twister3, Open Ephys) (https://open-ephys.org/twister3). We used tetrodes as the driving electrodes and tungsten wires (715000, A-M Systems, U.S.A.) as the fixed electrodes.

### Stereotaxic surgery for microdrive implantation

Surgical procedures for chronic electrophysiological recordings were performed under sterile conditions. After the training sessions were completed, mice were anesthetized with a cocktail of ketamine (100 mg/kg) and xylazine (10 mg/kg). Body temperature was monitored and maintained at 37 °C on a heating pad during the surgery. The mice were subcutaneously injected with 1% lidocaine (Aspen, Japan), and a midline incision was made to expose the skull and sutures. A stainless-steel anchor screw was implanted in the skull above the olfactory bulb, the skull surface was covered with dental adhesive resin cement (Super-Bond, Sun Medical, Japan), and the reference and grounding screw electrodes were embedded in the skull above the occipital cortex. Teflon-coated silver wires (786000, A-M systems, U.S.A.) were placed under the skin as electromyogram (EMG) unipolar electrodes. Tiny cranial windows were created above the bilateral motor cortical region through a cranial window (AP: +0.5 mm, ML: 1.2–1.5 mm to Bregma). As previously mentioned, a custom-made microdrive was used to insert the tetrodes and tungsten wires within the cranial windows. To secure the tetrodes from fixing with dental resin, grease was placed around the tetrodes. The microdrive was fixed with a dental resin (Unifast II, GC, Japan). After recovery for at least one week, food intake in the home cages was regulated to utilize a rodent tablet as a reward for the execution of reaching behavior, although water was available *ad libitum* in the cage. Food was supplemented every weekend.

### Forelimb reaching task

Mice used for skilled behavioral experiments regulated food intake but not water intake to allow them to engage in behavioral tasks. The body weights of these mice were monitored daily and maintained at 80–90% of their free-feeding body weight before behavioral training. Mice were placed in a custom-made transparent acrylic chamber (200 mm tall, 195 mm deep, and 90 or 140 mm wide, measured from the outside; 4 mm thick clear acrylic plate) and trained to perform the single-tablet retrieval task (Farr and Whishaw 2002). Mice learned to reach a 10-mg rodent tablet (5TUL, TestDiet, U.S.A.), which was manually placed on a platform outside the chamber. The chamber contained 9-mm-wide vertical openings at the middle of the chamber for mice to reach through with either the left or right paw. The platform (8.5 cm long, 4 cm wide, and 1.2 cm tall) contained a dip that was at a distance of 8 mm away from the edge of opening. For the unilateral paw-reaching task, the dip position was chosen to make the mice predominantly use the left paw. For the bilateral paw-reaching task, the dip position was adjusted to allow the mice to use either the left or right paw for reaching. During the initial shaping, the dip position was set in front of the opening at a distance to allow mice to reach the tablet with either paw (reaching), but not with the tongue (licking). Once the mice learned to reach the tablet, the dip position was displaced far from the opening, and the mice were allowed to use either of the paws to reach. Once the mice successfully reached and retrieved the tablet 10 times, the training was started. During the training session, we assessed the behavioral performance of the reaching-to-grasp task by counting the number of reaching attempts that touched a tablet placed on the dip. It was determined whether the tablet was moved by the attempt or retrieved inside the chamber. A single training session consisted of 120 attempts (30 min per day). For the bilateral paw reaching task, the dip position was switched when the mice revealed 20 reaching attempts using the demanded paw. Behavioral performance was assessed by measuring the rate of successful retrieval per session. The training sessions lasted until the mice reached a criterion (higher success rate reached 40% and lasted at least two consecutive daily sessions after five training sessions) or until the training sessions were over 12 sessions, even if the performance did not reach the criterion.

### Electrophysiological recordings from freely moving mouse

After the training sessions, the mice underwent the stereotaxic surgery described above to implant a microdrive that allowed the tetrode and tungsten electrodes to be inserted into the bilateral motor cortices. The mice were manually handled for 5 min over various days after a 1-week recovery period. After 5 days of retraining sessions, electrophysiological recordings were conducted during the reaching task. Mice performed at least 240 reaching attempts using the rodent tablet. Additionally, recordings were taken during a continued period lasting 10–15 min when mice were not provided with the rodent tablet and, thus, were not engaged in reaching behavior. Multiunit and EMG signals were amplified and digitized by a headstage (C3314, Intan Technologies, U.S.A.), sampled at 20 kHz with an interface (C3100, Intan Technologies, U.S.A.), and acquired with a personal computer. Videos were recorded at a 60 Hz frame rate and 720 × 480 pixels (GV-SDREC, I-O data device, Inc., Japan). To synchronize physiological data with video images in *post hoc* offline analysis, a synchrony signal generated by an Arduino microcontroller was recorded by the data acquisition interface for physiological data and video through an audio input channel, as demonstrated previously (Topper et al. 2014). After the end of recording sessions, electrolytic lesions were made through tetrodes by passing continuous DC current (+2.5–3.0 µA for 10 s) using an isolator (ISO-Flex, A.M.P.I., Israel).

### Electrophysiological recordings under anesthetized mouse with pharmacological manipulation

For electrophysiological analyses of interhemispheric interactions, mice were anesthetized with urethane (1.5 g/kg, i.p.). The animal was fixed in a stereotaxic frame, and a midline incision was made to expose the skull surface. Reference and ground-screw electrodes were implanted in the skull over the cerebellum. In the case of topical drug application to recording site, a cranial window was made and dura matter was kept intact at rostral motor cortical area (AP: +2.0 mm, ML: 0.5 mm from Bregma) of each hemisphere. A “well” surrounding the cranial window at left hemisphere was built with dental adhesive resin cement (Super-bond, Sun medical, Japan) to enable topical application of drugs over the recording site. To acquire local field potentials (LFPs) along the cortical laminar structure, a 16-ch silicon probe (A1×16-5mm-150-177-A16, NeuroNexus Technologies, U.S.A.), in which each electrode was spaced vertically (0.15 mm, was vertically inserted within the left cranial window to allow the top electrode to be placed in layer 1. A tetrode array consisting of two tetrodes was inserted in the same cranial window. The silicon probe was connected to a headstage adaptor (A16-OM32, NeuroNexus, U.S.A.) installed on a micromanipulator (SM-25A, Narishige, Japan) mounted on a stereotactic frame (SR-9M, Narishige, Japan). A headstage (C3314, Intan Technologies, U.S.A.) was connected to the adaptor. For tetrode recordings, the other end of the exposed tetrode wire was connected to a normal-pitch gold pin using electrically conductive pasts (Dotite, Fujikura Kasei Co., Ltd., Japan). A 36-pin wire adapter (C3313, Intan Technologies, U.S.A.) was connected to the headstage (C3314, Intan Technologies, U.S.A.), and the wires of this adapter were connected to gold-pin connectors. The tip of a bared optical fiber (M37L-02, diameter = 0.55 mm, 0.22 NA, Thorlabs, U.S.A.) was placed at 0.2 mm above the right cranial window and was covered 1.2% agar and paraffin to prevent from dry-out. We recorded LFPs and multiunit activity for at least 30 min as a control condition while phosphate buffer saline (PBS, 0.1 M, 3–4 µl) was filled in the well. We replaced the solution with either of cholinergic agonist (50 µM, 3–4 µl), carbachol (Sigma-Aldrich), or muscarinic acetylcholine (ACh) receptor antagonist, scopolamine (22 mM, 3-4 µl, Tokyo chemical industry). After the first drug condition, the solution in each well was replaced with PBS as a wash condition. After washing for 20 min, the solution was replaced with another drug solution. The order of drug application was randomly chosen for each experiment.

In another case of systemic administration of drug, a cranial window was made and dura matter was kept intact at motor cortical area (AP: +0.5–1.0 mm, ML: 1.2 mm from Bregma) of each hemisphere. To acquire multiunit activity and LFPs from the bilateral motor cortex, a tetrode array consisting of four tetrodes was inserted within each cranial window to allow the tetrode array to cover 0–+1.0 mm on the AP axis. The tetrode wires were connected to the headstage as described above. We recorded multiunit activity for 30 min as a control condition and intraperitoneally injected either of rivastigmine (3.6–4.0 µM/kg, Sigma-Aldrich, U.S.A.) or scopolamine (0.5 mg/kg). We continued recording for 60 min. Multiunit signals were amplified and digitized using headstages (C3314, Intan Technologies, U.S.A.), sampled at 20 kHz with an interface (C3100 or C3004, Intan Technologies, U.S.A.), and acquired using a personal computer.

### Optogenetic stimulation

Optogenetic stimulation was delivered by using a LED light source (Optogenetics-LED-Blue, 460 nm, Prizmatix Ltd., Israel) triggered by 1-ms TTL single pulse generated with a TTL pulser (StimJim, Open Ephys, https://open-ephys.org/stimjim) controlled by Arduino. The light intensity was controlled to set the maximum emission power approximately 2– 10 mW (8.4–42 mW/mm^2^ at cortical surface), which is within a safe range for *in vivo* experiment (about 75 mW/mm^2^ for pulse duration ranging between 0.5 and 50 ms) as shown in previous publication (Cardin et al. 2010). A single pulse stimulation was repeated 40 times with various inter-stimulus intervals in the range of 10–11 s before the application of the first drug and at 5 min and 15 min after the application of each drug.

### Histology

For immunohistochemistry, mice were deeply anesthetized with an overdose injection of urethane and transcardially perfused with 0.1 M PBS followed by 4% paraformaldehyde (PFA) in 0.1M PBS (pH 7.4). After post-fixation in PFA overnight, 75-µm-thick sections (150 µm apart) were cut using a vibratome (DTK-1000N, D.S.K, Japan) and processed for immunohistochemistry. After the sections were washed in 0.1 M PBS, they were incubated for 1 h at room temperature (20– 25 °C) in 0.1 M PBS with 1% blocking reagent (Roche) and 0.3% Tween 20 (Sigma-Aldrich). Primary antibodies were diluted in 0.1 M PBS buffer containing 1% blocking reagent and 0.3% Tween-20. Sections were incubated with monoclonal rabbit anti-parvalbumin (PV) antibody (1:500, ab181086, Abcam) and rat anti-muscarinic ACh receptor m2 (M2R) antibody (1:500; MAB367, Sigma-Aldrich) at 4 °C for 24 h on a shaker. After sections were washed in 0.1 M PBS, they were incubated with secondary antibodies, including donkey anti-rabbit IgG-Alexa Fluor 488 (1:500, ab150073, Abcam) and goat anti-rat IgG-Alexa Fluor 594 (1:500, 405422, BioLegend) at 4 °C for 24 h on a shaker for fluorescent immunostaining. After the secondary antibody reactions, sections were counterstained with DAPI. Fluorescence imaging was performed using an epifluorescence microscope (MVX10, Olympus, Japan) and a confocal spinning disk microscope (CSU-X with iXon Ultra897, Andor Technology Ltd., UK). Z-stack confocal imaging was performed over unilateral or bilateral motor cortical areas (AP: 0–+1.95 mm, ML: 1–2 mm), in which all cortical layers were scanned by shifting the field of view along the cortical depth from the pia matter to the white matter. The coordinates of the motor cortical regions are in accordance with those of the mouse brain atlas (Franklin and Paxinos, 2012).

### Data analysis

All behavioral and neuronal data were analyzed by custom-written MATLAB scripts (The MathWorks, Inc.) or by an open source Python program.

#### Behavioral analysis

The paw movements of the mice filmed by a camera placed at the side of the chamber, as described above, were processed using a marker-less pose estimation algorithm, DeepLabCut (Mathis et al. 2018) (https://github.com/AlexEMG/DeepLabCut), which allowed us to estimate how the mouse moved its forepaws to reach and grasp the rodent tablet. To detect the trajectory of each paw during reaching, we analyzed the temporal sequence of paw positions on the horizontal axis of the video, which enabled us to detect the forward and backward motion of the paw during reaching after the paw exited through the vertical opening of the chamber. We removed the trajectory of the unmoved paws using the empirically set threshold speed of the trajectory. The maximum position of each trajectory is defined as the endpoint of the reaching movement. We used reaching trajectories for further analyses of electrophysiological data, regardless of the presence or absence of rodent tablets on the platform.

#### Spike sorting, clustering, and classification of putative cell-types

Spike events from multiunit activity were isolated using a custom-made semi-automatic spike-sorting program, EToS [12 feature dimensions for four channels; high-pass filter, 300 Hz; time resolution, 20 kHz; spike-detection interval, >0.5 ms) (Takekawa et al. 2010, 2012), as done previously (Handa et al. 2017, 2021). The sorted spike clusters were combined, divided, and manually discarded to select single neuron clusters using Klusters (Hazan et al. 2006). After manual clustering, we classified putative cell types based on two parameters: spike width (trough-to-peak) and burstiness of spiking (rise time of autocorrelogram [ACG]) using CellExplorer (Petersen et al. 2021) (https://cellexplorer.org/). Units revealing narrow spike width (≤0.45 ms) were classified as putative fast spiking (FS) interneurons. Other units revealing wider spike width (>0.45 ms) were classified based on the rise time of ACG into wide interneuron (>6 ms) and putative regular spiking (RS) neurons (≤6 ms).

#### Reaching-related neuronal activity

Reaching-related neuronal activity was examined in well-isolated units. To make peri-event time histograms (PETHs), we calculated the mean instantaneous firing rate in a 20-ms bin around endpoint of reaching in a range from −1 s to +1 s. To determine if the unit activity was modulated by reaching, firing rates at two periods of 0.2 s before and after endpoint of reaching were compared with control activity, which was mean firing rate for 0.4 s from −1 s relative to time at the endpoint of reaching, by two-sample Kolmogorov–Smirnov (KS) test (*P* <0.05) with Bonferroni correction. If there were statistically significant differences in the firing rates in either period, we defined the unit as a reaching-related neuron. To classify reaching-related activity, the PETHs were smoothed with a Gaussian filter (s = 20 ms) and normalized by subtracting the mean of the control activity and dividing by the standard deviation (SD) of the control activity. If the peak value of the normalized PETH level was positive (or negative), the neuron was defined as Up- or Down-activity type. To quantify the relevance of neuronal activity to reaching behavior, we calculated the KS statistic *K* as follows:

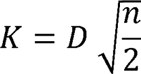

Here, *D* and *n* indicate the KS test statistic and number of attempts, respectively. To profile the firing pattern of collective neurons across multiple recording sessions and mice (nine sessions, four mice) during the unilateral paw-reaching task, the z-scored PETHs were further normalized by the absolute peak value. To classify the characteristics of neuronal activity during the bilateral paw reaching task (six sessions, two mice), we computed the PETH for each paw reaching task and assessed whether the activity was related to reaching using the two-sample KS test (*P* <0.05) with Bonferroni correction, as described above. For significant neurons, we determined the excitatory or inhibitory activity type in each paw (ipsilateral or contralateral), as described above.

#### Interhemispheric spike-triggered average (STA) analysis

To examine the interhemispheric spike correlation during the reaching task, the spike-triggered average (STA) was computed under two conditions: when mice performed the unilateral-paw reaching task and when they were not engaged in performing the task due to the absence of a rodent tablet. We used spike trains of contralateral motor cortical neuron as a reference time and computed the number of spikes of ipsilateral motor cortical neuron in a window ranging from −0.5 s to +0.5 s aligned at each spike time of the contralateral motor cortical neuron (bin = 5 ms). Significance of spike synchrony (or positive correlation) was detected by comparing the positive peak value of the STA with the mean+4SD of values within a baseline window (−0.5 to −0.2 s and +0.2 to +0.5 s) under the task condition. If the peak value of a cell pair was larger than this criterion, we judged that the cell pair showed synchronous excitatory firing activity between the ipsilateral and contralateral motor cortices. In contrast, the significance of synchronous suppression (or negative correlation) was detected by comparing the negative peak value in the computed STA with the mean-4SD of values in a baseline window. If the peak value of a cell pair was smaller than this criterion, we judged that the cell pair showed synchronous inhibition of the ipsilateral cortical neurons. To investigate the pharmacological effect on spike synchrony under anesthesia, we computed the STA between left motor cortical neurons and right motor cortical neurons at four time points (control for 15 min before drug administration, three consecutive 15-min periods after drug administration), and the significant synchronous activity was judged with the same criterion as described above.

#### Analysis of neural activity evoked by a brief optogenetic stimulation

To investigate the profiles of LFPs in response to brief optogenetic stimulation at the contralateral motor cortex, the acquired LFPs were downsampled at 1 kHz, followed by band-pass filtering (3–300 Hz). For the linear probe data, we analyzed the LFPs recorded at the top nine electrode positions covering the cortical layers 1 to 6 (Fig. 5A). The evoked LFPs were averaged over 40 optogenetic stimulation trials conducted during the control period and 15 min after topical application of each drug. A common response pattern of evoked LFP was an early large negative deflection followed by a positive deflection, as observed in Fig. 5B and previously reported (Spalletti et al. 2017). To quantify the profiles of the trial-averaged LFP responses, we measured the area above the largest negative deflection, which emerged within 50 ms after stimulation. The area under the large positive deflection that emerged within 350 ms after stimulation was computed. For each property, we assessed the difference between electrode positions (i.e., electrode depth) and drug conditions using two-way ANOVA. If the difference in drug conditions was statistically significant (*P* <0.05), we further assessed the difference between the control and each drug condition at each electrode depth using a paired t-test with Bonferroni correction (*P* <0.05). For the LFP and multiunit activity recorded with the tetrodes, we analyzed the LFP data using the same process as that for the linear probe data, whereas we computed the peri-stimulus time histogram in response to optogenetic stimulation using sorted spike activity.

#### Analysis of RNA-seq data

Previously published single cell RNA-seq data (a data set “Primary Motor Cortex from Mouse - Single Cell 10X RNAseq -V3”) was re-analyzed to examine differential gene expression in subtypes of neurons in the murine primary motor cortex on Neuroscience Multi-Omic Analytics (NeMO) platform (Orvis et al. 2021; Yao et al. 2021) (https://nemoanalytics.org/). Based on the clustering of cell subtypes by Yao et al. (2021), plots were drawn for mRNA expression levels in representative excitatory (L5 IT Pld5, L5 ET, and L6 CT) and inhibitory neuronal clusters (Lamp5 Lhx6, Pvalb Gabrg1, Sst chodl, and Vip Htr1f) using the “Multigene Display Viewer” on the platform. We selected genes encoding the vesicular glutamate transporter, VGLUT1 (*Slc17a17*), GABAergic interneurons (*Gad1*, *Pvalb, Sst*, and *Vip*), muscarinic ACh receptors (*Chrm1*, *Chrm2*, *Chrm3*, *Chrm4*, and *Chrm5*), and nicotinic ACh receptors (*Chrna1*, *Chrna2*, *Chrna3*, *Chrna4*, *Chrna7*, *Chrna9*, *Chrna10*, *Chrnb1*, *Chrnb2*, *Chrnb3*, and *Chrnb4*).

#### Image processing and cell count

Acquired confocal fluorescence images were analyzed *post hoc* by means of ImageJ Fiji, an open-source platform for image analysis (Schindelin et al. 2012). We manually counted M2R-positive cells in each image and checked whether the cell was PV-positive cell or not across cortical layers in a range from layer 1 to layer 6. The data were acquired from five hemispheres from three mice.

## Results

### Reaching-related neuronal activity in the contralateral motor cortex is more relevant for reaching behavior than that in ipsilateral motor cortex

The mice were trained to reach for their left forepaw with a rodent tablet placed on a platform through the opening of the chamber (Fig. 1A). The trajectory of forepaw reaching revealed forward and backward motions and the peak position represented the endpoint of reaching (Fig. 1B). We recorded multiunit activity and LFPs from the bilateral motor cortical areas during the unilateral paw-reaching task (nine sessions from four mice). We recorded 192 single-unit activities in total and classified those units into putative cell types: RS cells, FS interneurons, and wide interneurons based on spike width and auto-correlogram tau rise (Fig. 1C), which indicate the physiological properties of neuron types, as described previously (Petersen et al. 2021). Most neurons (N = 184, 95.8%) exhibited reaching-related activity. We found that reaching-related activity showed an increase and/or decrease in the firing rate around the reaching end in both the ipsilateral and contralateral motor cortices (Fig. 1D). To investigate how different these reaching-related neuronal activities are between ipsilateral and contralateral motor cortical neurons and between cell types, we analyzed the activity pattern of individual neurons by classifying them into two cell types (RS cells and INT including FS and wide interneurons) and a firing pattern in each hemisphere (Up and Down type, showing increased and decreased activity upon reaching, respectively). Importantly, a substantial number of contralateral RS and ipsilateral INT cells showed Up type activity, whereas ipsilateral RS cells showed Down type activity (Fig. 1D). Furthermore, we quantified the relevance of neuronal activity to reaching behavior by calculating the relevant Kolmogorov–Smirnov statistics. In the contralateral motor cortex, the reaching relevance of RS-Up type neurons was greater than that of INT-Up type neurons (Mann–Whitney U test, *P* <0.01), whereas the reaching relevance of INT-Down type neurons was greater than that of RS-Down type neurons (Mann–Whitney U test, *P* <0.001) (Fig. 1E). In contrast, in the ipsilateral motor cortex, reaching relevance was not significantly different between cell types for any activity type (Mann–Whitney U test, Up-type: *P* = 0.291, Down-type: *P* = 0.745).

**Figure 1.**
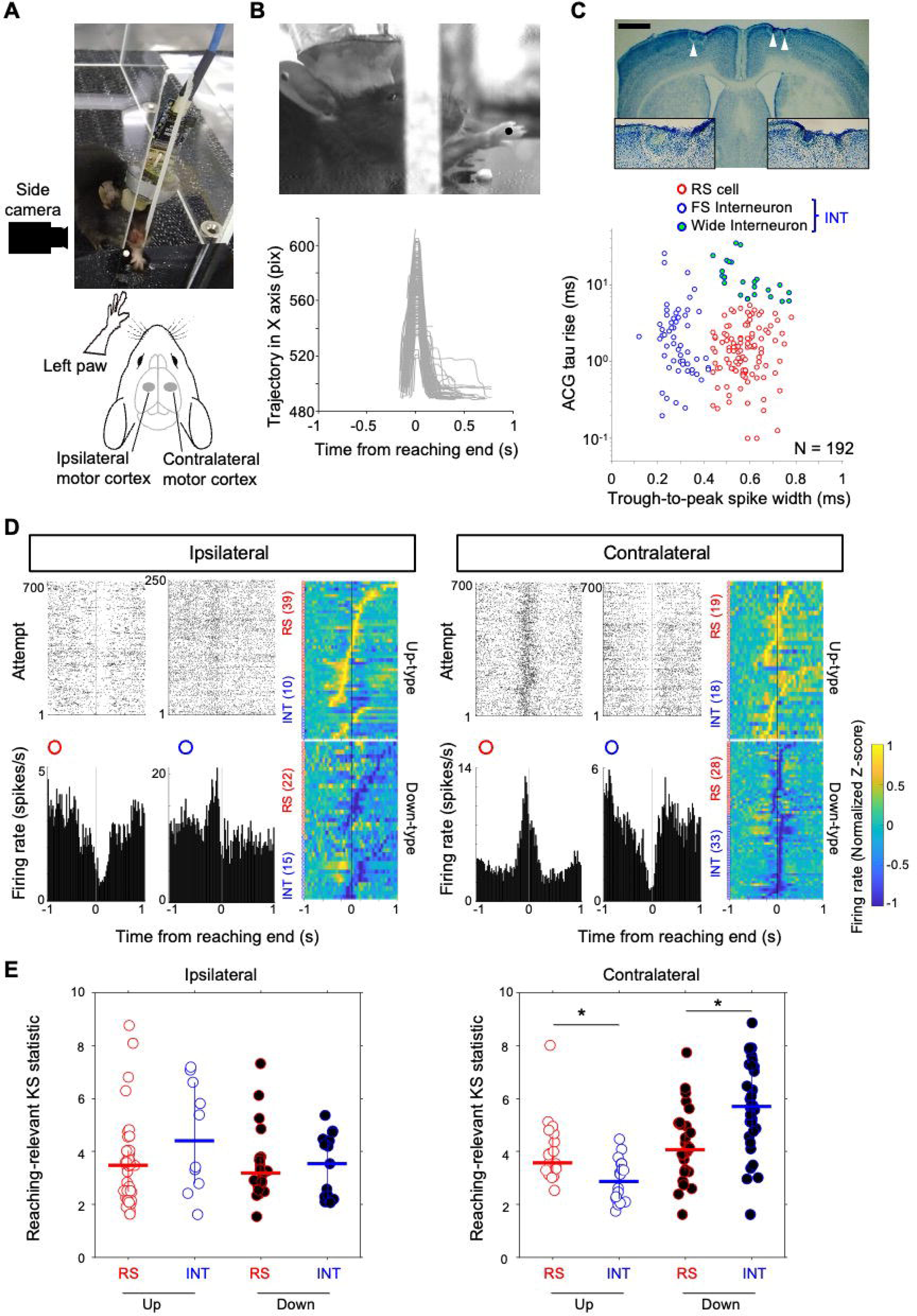
Reaching-related neuronal activity in ipsilateral and contralateral motor cortex. **(A)** Top: A snapshot of a mouse during reaching behavior to grasp a small tablet with its left paw. The animal placed in a transparent chamber with a slit through which the mouse reaches its paw. A microdrive implanted on the skull is connected to a headstage with digital cable (blue). A camera is set at the right side of chamber. Bottom: A schematic illustration of a left paw for reaching and the recording locations in the bilateral motor cortices. **(B)** Top: A snapshot recorded by a side camera. A mouse reached its left paw toward a tablet placed at a stage set in front of the slit of the chamber. A black dot indicates an instantaneous paw position. Bottom: Representative trajectories of left paw during reaching along horizontal axis. The time series of trajectories are aligned at the endpoint of reaching. **(C)** Top: Locations of micro electrolytic lesions made after recording in a Nissl-stained brain coronal section (white arrowheads). Scale bar, 1 mm. Insets, magnified views of area around lesions indicated by white arrowheads in left and right hemispheres. Bottom: Classification of putative cell-types based on the spike width and rise time in auto-correlogram (ACG). **(D)** Representative firing patterns of the putative regular spiking (RS) and interneuronal (INT) cells in the ipsilateral (left) and contralateral cortices (right) during reaching. Collective firing activity is presented in heatmap for ipsilateral and contralateral cells by separating putative cell-types with firing patterns: Up- (top) and Down-types (bottom). **(E)** Quantitative comparison of reaching-relevant KS statistics between RS and INT cells for each firing pattern in ipsilateral (left) and contralateral (right) motor cortices. Horizontal and vertical lines present median and the range between the first and third quantiles, respectively. Asterisk indicates a statistical significance assessed by Mann–Whitney U test (*P* <0.01).

These results suggest that the contralateral RS-Up and INT-Down type neurons are more relevant to the reaching task and that these neurons could contribute to excitation in the contralateral motor cortex, and thus to interhemispheric inhibition via the ipsilateral INT-Up neurons to inhibit the ipsilateral motor cortex.

### Reaching-related activity is significantly modulated by paw-laterality

Next, we examined whether reaching-related activity was modulated by the laterality of the paw (i.e., contralateral and ipsilateral paws) using the unilateral paw reaching task. If reaching-related neurons, especially the contralateral RS-Up and INT-Down types, contribute to the excitability of the contralateral motor cortex depending on the contralateral paw, then reaching-related activity should be altered by changes in paw laterality. To address this question, we recorded multiunit activity from the bilateral motor cortical areas during a bilateral paw-reaching task in six sessions in two mice (Fig. 2A). Of the 92 reaching-related neurons detected during the task with forced contralateral paw use, 43 and 48 were classified as putative RS and INT cells, respectively. We observed that Up-type neurons activated upon contralateral paw use responded less and were even inactivated by ipsilateral paw use, irrespective of the cell type (Fig. 2B). This was also the case for Down-type neurons with inactivation upon contralateral paw use, showing a lower response to ipsilateral paw use (Fig. 2C). Population analysis revealed that the peak firing rate at the time of reaching both Up- and Down-type neurons was significantly modulated by paw laterality in both cell types (Wilcoxon signed-rank test, RS-Up type N = 24, *P* <0.001, INT-Up type N = 24, *P* <0.01, RS-Down type N = 19, *P* <0.01, and INT-Down type N = 24, *P* <0.05) (Fig. 2D).

**Figure 2.**
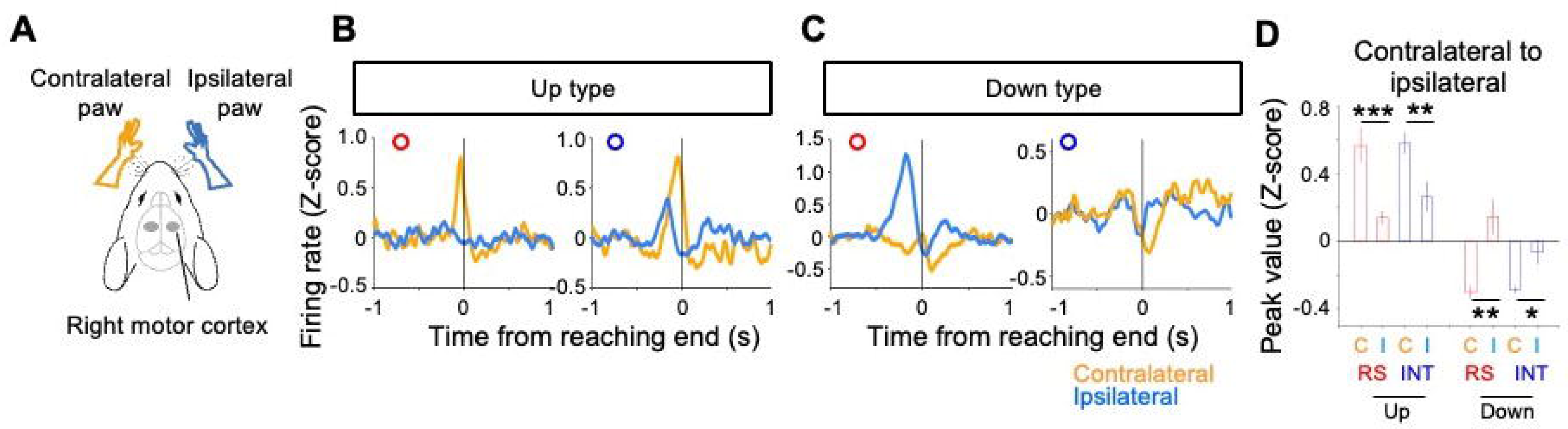
Reaching-related activity is dependent on paw-laterality. **(A)** A schematic illustration to represent a relationship between neurons of interest and contralateral (orange) and ipsilateral paws (blue) in case those neurons were recorded in the right motor cortex. Otherwise, *vice versa*. Gray shaded area indicates recording site. **(B–C)** Four representative neuronal activity during reaching for Up-type (**B**) and Down-type (**C**) firing pattern of each cell-type (RS and INT). The firing pattern is judged by firing activity under contralateral condition. Red and blue circles indicate RS and INT cell-types, respectively. **(D)** Comparison of peak values of PETH when contralateral paw (**C**) and ipsilateral paw (**I**) conditions for each firing pattern and cell-type. Values are presented by mean and standard error of mean (SEM). Statistical significance is assessed by Wilcoxon signed rank test. *: *P* <0.05, **: *P* <0.01, ***: *P* <0.001.

This result suggests that reaching-related activity is strongly modulated by paw laterality and may reflect changes in the excitatory-inhibitory balance within the unilateral motor cortex depending on paw laterality for reaching behavior.

### Enhancement of interhemispheric spike correlation during reaching task

Excitability of the contralateral motor cortex during reaching may affect neuronal activity in the ipsilateral motor cortex through interhemispheric connections. To address whether the interhemispheric interaction emerges during reaching behavior, we analyzed spike-triggered average (STA) between ipsilateral and contralateral motor cortical neurons (N = 461 cell pairs) during unilateral-paw reaching task performance, as well as during the non-task period when the mice were not engaged to perform the task by quitting delivery of the rodent tablet (Fig. 3A). We found that a substantial number of interhemispheric neuronal pairs covaried positively (yellow in Fig. 3C) and negatively (blue in Fig. 3C), as revealed by the higher peak value in the STA during the reaching task compared with the period without the task (Fig. 3B). More specifically, we found an enhancement of the negative correlation in STA between ipsilateral and contralateral RS cells (right in Fig. 3D), with statistically significant changes in STA peak amplitude (right in Fig. 3F; Wilcoxon signed rank test, N = 6 pairs, *P* <0.05), but no significant change in the positive correlation in STA between ipsilateral and contralateral RS cells (left in Fig. 3D and 3F; Wilcoxon signed rank test, N = 8 pairs, *P* = 0.148). In contrast, we found an enhancement of the positive correlation in STA between ipsilateral FS neurons and contralateral RS neurons (left in Fig. 3E) with statistically significant changes in STA peak amplitude (left in Fig. 3G; Wilcoxon signed-rank test, N = 11 pairs, *P* <0.05), but no significant change in the negative correlation in STA between ipsilateral FS neurons and contralateral RS neurons (Wilcoxon signed-rank test, N = 6 pairs, *P* = 0.0625) (right in Fig. 3E and 3G).

**Figure 3.**
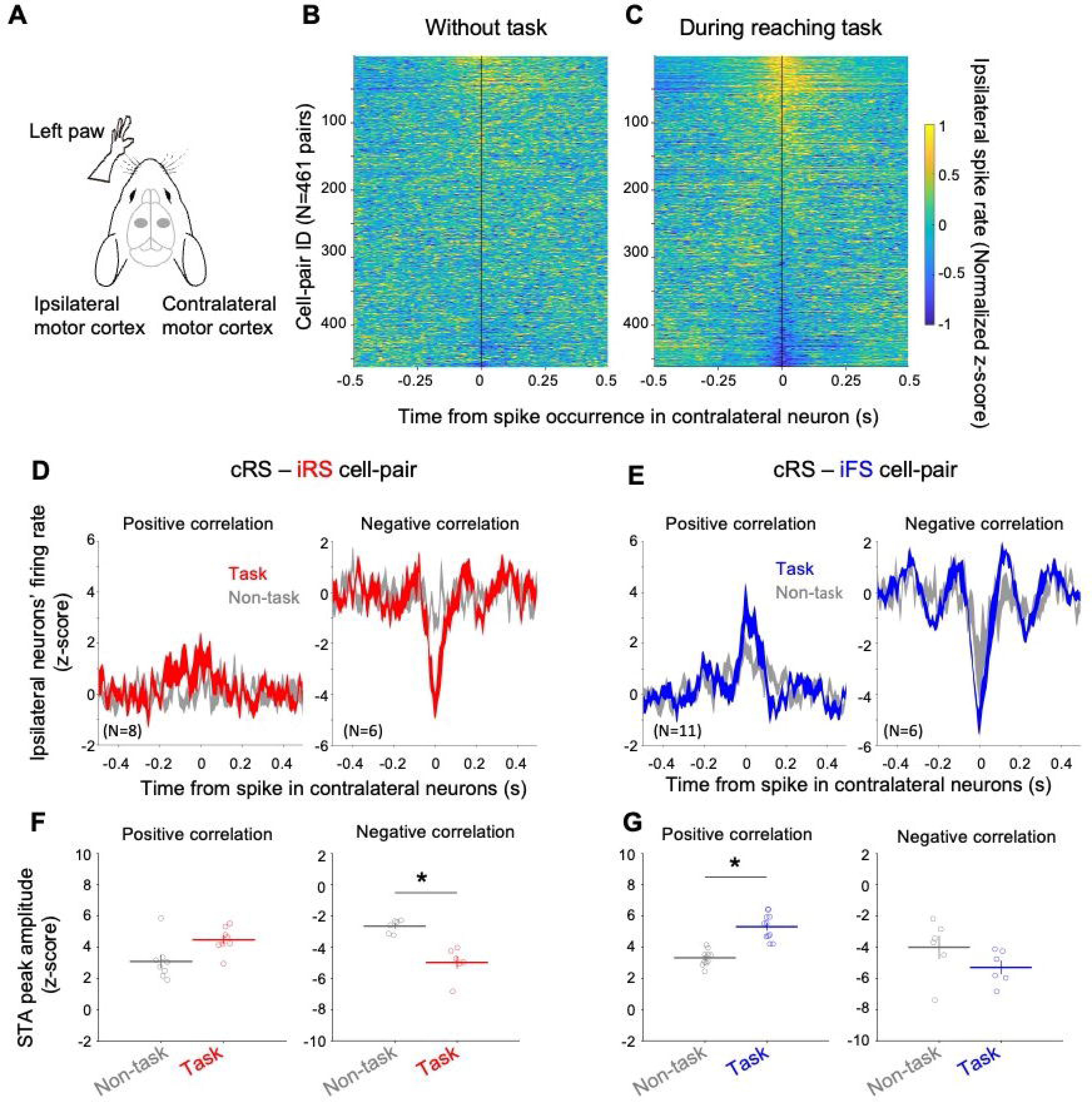
Enhancement of interhemispheric spike correlation during reaching task. **(A)** A schematic illustration of a left paw for reaching and the recording locations in the bilateral motor cortices. **(B–C)** Spike-triggered average (STA) between ipsilateral and contralateral cortical neurons without (**B**) and during reaching task (**C**). The heatmap shows normalized spike count of ipsilateral neuron relative to time of spike events in the contralateral cortical neuron recorded in the same recording session. **(D–E)**: Collective STA data from two cell-pairs, contralateral RS and ipsilateral RS (cRS-iRS) cell-pair (**D**) and contralateral RS and ipsilateral FS (cRS-iFS) cell-pair (**E**). The STAs in red or blue show positive (left) and negative (right) correlation in each cell-pair during task. The STAs in gray indicate the STAs without task by the same cell-pairs. **(F–G)** Comparison of peak values of STA for those cRS-iRS cell pairs (**F**) and cRS-iFS cell pairs (**G**) between recording periods with (colored) and without task (gray). Horizontal and vertical lines indicate mean and SEM, respectively. An asterisk indicates statistical significance by Wilcoxon signed rank test, *P* <0.05).

This result suggests that the interhemispheric interaction is enhanced by the reaching task, and the negative correlation between RS and RS and the positive correlation between RS and FS may reflect the involvement of interhemispheric inhibition in suppressing the excitability of the ipsilateral motor cortex.

### Cholinergic modulation of interhemispheric spike correlation

To address the role of ACh in the modulation of the interhemispheric synchronous spike correlation observed during task performance, we used tetrode arrays to examine the effect of pharmacological cholinergic modulation on ongoing neuronal activity in the bilateral motor cortices of anesthetized mice (N = 3 mice for each drug condition) (Fig. 4A). To monitor the ongoing activity of neurons in both hemispheres, we systemically administered the cholinesterase inhibitor, rivastigmine, or the muscarinic receptor antagonist, scopolamine. STA analysis revealed that rivastigmine impaired the positive correlation with reduced peak amplitude in STA compared to the control (paired t-test: *P* <10^-120^, Fig. 4B). In contrast, scopolamine enhanced the positive correlation with a significant increase in peak amplitude in the STA compared to that in the control (paired t-test: *P* <10^-17^, Fig. 4C). The change in the peak amplitude in STA showed a statistically significant contrast between the drug conditions (two-sample t-test with Holm–Bonferroni correction, *P* <10^-101^) (Fig. 4D). We could not attribute these changes to the altered firing rate under drug treatment, since the ongoing firing activity of RS and FS neurons after the systemic administration of rivastigmine (paired t-test: RS neurons, N = 240, *P* = 0.825; FS neurons, N = 34, *P* = 0.0912) and scopolamine (paired t-test: RS neurons, N = 216, *P* = 0.614; FS neurons, N = 29, *P* = 0.974) did not significantly change during the recording period as compared with the control period.

**Figure 4.**
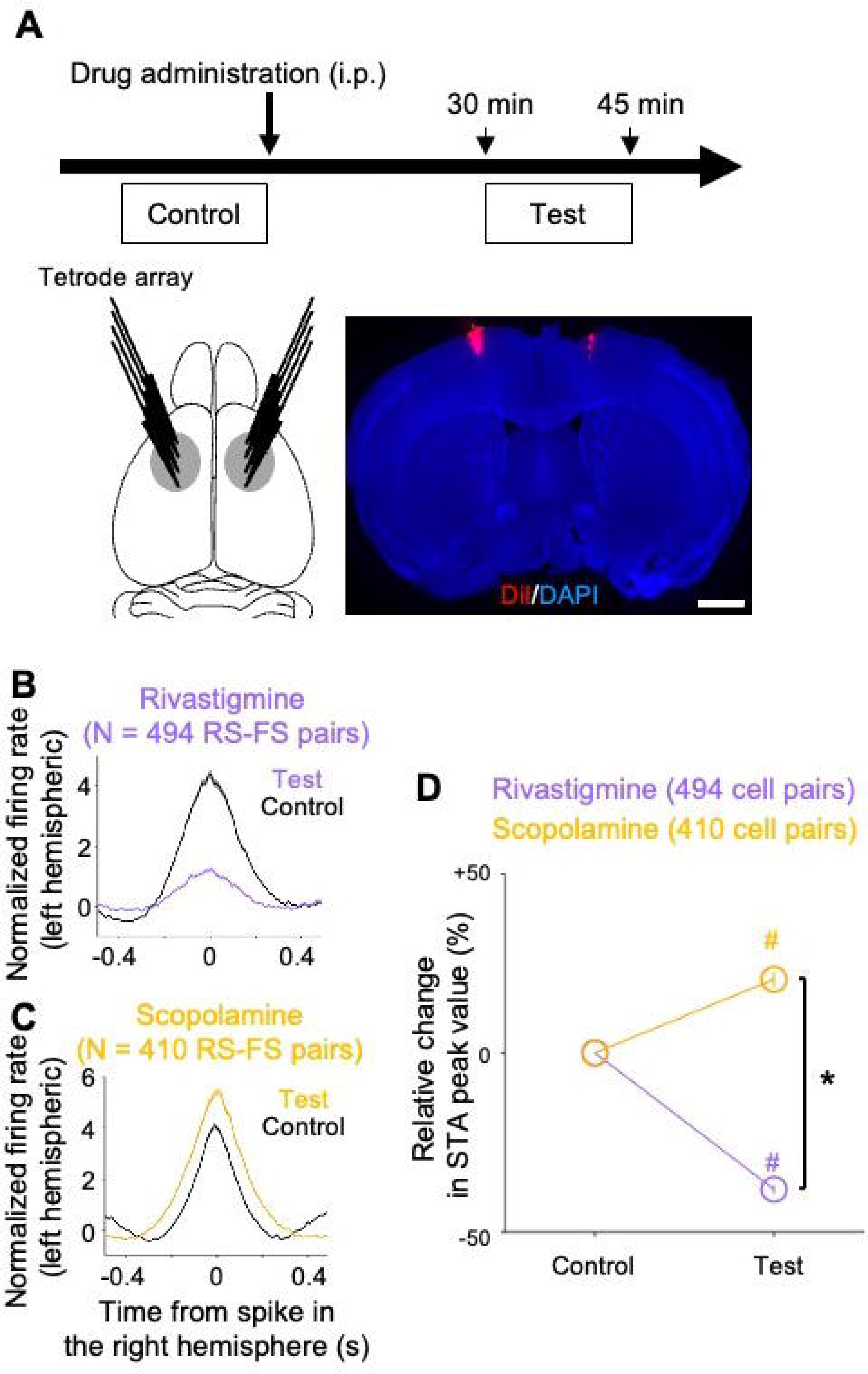
Cholinergic modulation of interhemispheric spike correlation. **(A)** Top: A schematic showing an experimental design. Analysis periods are shown as a control period for 15 min before drug administration and a test period for 15 min from 30 min after drug administration. Bottom left: A schematic illustration of multiunit recording sites in the bilateral motor cortices with tetrode arrays. Bottom right: A coronal section revealing recording sites based on the track of tetrodes coated with DiI in the bilateral motor cortices. Scale, 1 mm. **(B)** Comparison of STA of RS-FS cell-pairs between control and test periods. Shaded area shows SEM. **(C)** Comparison of change in peak value of STA relative to control in test period (paired t-test, #: *P* <0.0001) and between drug conditions (two-sample t-test with Holm–Bonferroni correction, *: *P* <0.0001). Values are presented by mean and SEM.

These results suggest that the interhemispheric interactions between RS and FS neurons are differentially modulated by ACh efficacy.

### Cholinergic modulation of interhemispheric inhibition in deep layers

We addressed how interhemispheric inhibition through callosal inputs was affected by cholinergic modulation. We recorded the LFPs along cortical layers in response to the photostimulation of the contralateral motor cortex using Emx-1-Cre;Rosa-ChR2-EYFP mouse expressing ChR2-EYFP in the entire cerebral cortex (Fig. 5A). We placed a linear silicon probe with 16 electrodes covering the entire extent of the six cortical layers in the left motor cortex (Fig. 5A). A common response pattern of the evoked LFP revealed a large negative deflection followed by a positive deflection (top in Fig. 5B). Simultaneous recording of the spike activity of RS neurons revealed a brief excitation during the negative deflection of the evoked LFP, followed by long-lasting suppression during the positive deflection of the evoked LFP (bottom in Fig. 5B). After recording the evoked response to brief optogenetic stimulation as a control, a solution of either the muscarinic agonist carbachol or antagonist scopolamine was topically applied to the recording site. The results showed that the area under the positive deflection was significantly enhanced by scopolamine, but not by carbachol, whereas the area above the early negative deflection did not change (Fig. 5C). Statistical analyses of the area under the deflections (Fig. 5D) revealed that scopolamine significantly enhanced the area under the positive deflection in the deeper layers corresponding to deep layers 5 and 6 (Fig. 5D1; two-way ANOVA, *P* < 0.05, *post hoc* paired t-test with Bonferroni correction, *P* <0.05), but not the area above the negative deflection (Fig. 5D2; two-way ANOVA, *P* = 0.471).

**Figure 5.**
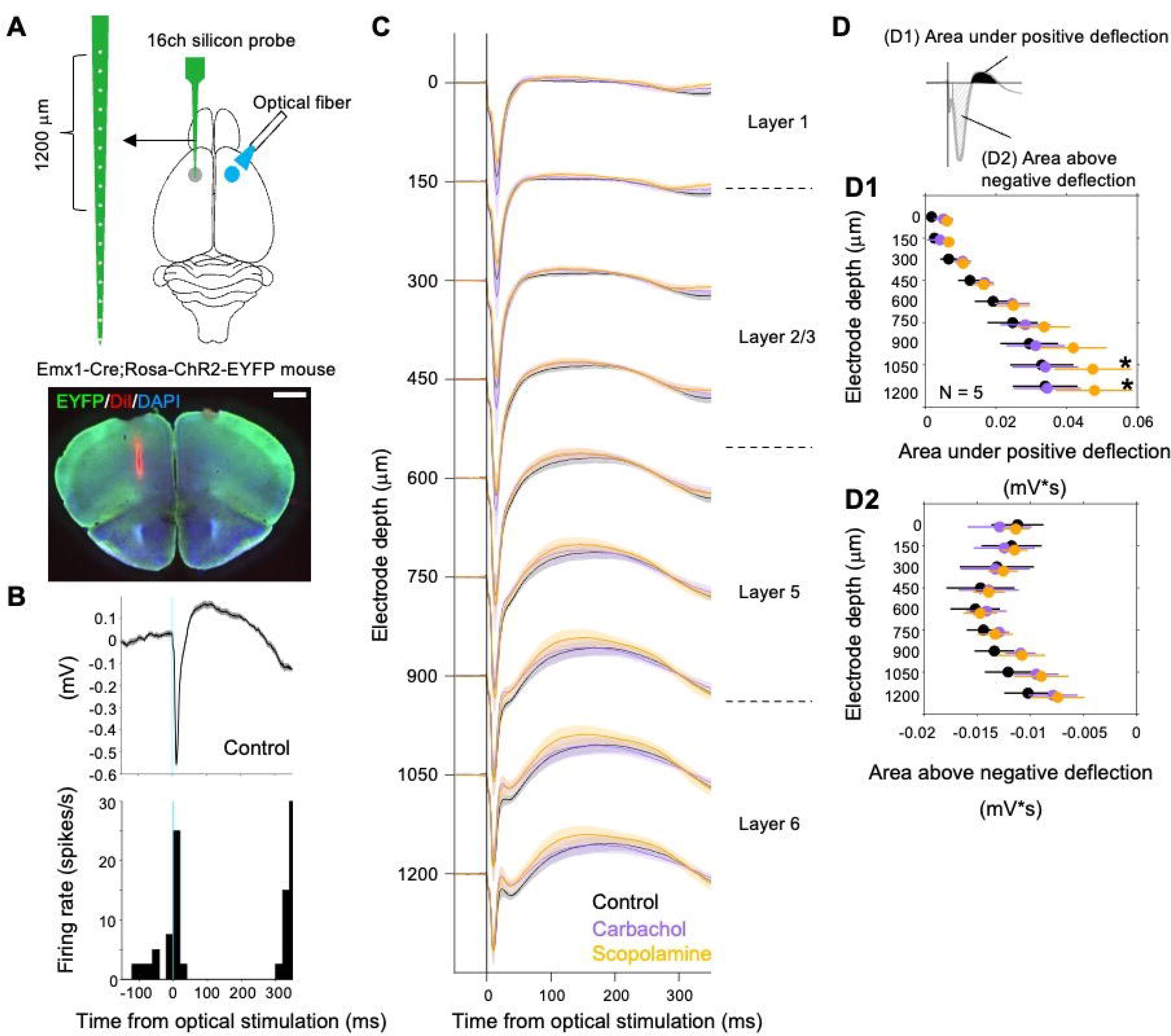
Cholinergic modulation of interhemispheric inhibition in deep layers. **(A)** An experimental design of electrophysiological recording in the left motor cortex of Emx1-Cre; Rosa-ChR2-EYFP mouse in response to optogenetic stimulation of the right motor cortex. A linear probe with 16-channel was coated with DiI and placed to the left motor cortex as shown by a coronal section shown on the right. Scale bar, 1 mm. **(B)** Top: Trial-averaged LFP in response to contralateral optogenetic excitation (light blue line). Bottom: An example of spike activity of a simultaneously recorded neuron. (**C**) Comparison of population LFP responses to contralateral optogenetic excitation among control (black), carbachol (purple), and scopolamine (orange) at each electrode depth. Solid line and shaded area represent mean ± SEM. Indicated cortical layer on the right is based on the atlas of mouse brain (Franklin and Paxinos, 2019). (**D**) Top: A schematic illustration of measured areas under positive deflection (black solid) and above negative deflection (hatched) of LFP responses. (**D1**, **D2**) Comparison of characteristics of evoked LFP response among drug conditions regarding area under positive deflection (**D1**) and area above negative deflection (**D2**). An asterisk indicates the statistical significance between control and scopolamine groups at each electrode depth (two-way ANOVA, *post hoc* paired t-test with Bonferroni correction, *P* <0.05). Values are presented as mean ± SEM.

These results suggest that blocking ACh binding to the muscarinic ACh receptor enhances cortical inhibition through neural responses to callosal inputs. Moreover, cholinergic modulation through muscarinic ACh receptors differentially influences interhemispheric inhibition along cortical depths, particularly in the deep layers (deep level of layer 5 and layer 6).

### Expression of muscarinic acetylcholine receptor in interneurons of deep layers of motor cortex

To explore the potential cellular elements that could contribute to the cholinergic modulation observed in deep layers, we analyzed the mRNA expression of ACh receptor subclasses in mouse motor cortical neurons using an open-source platform (Yao et al. 2021), which enabled us to analyze cell-type-based mRNA expression using publicly available single-cell RNA-seq datasets. We found that three gene markers of muscarinic ACh receptors, *Chrm1*, *Chrm2*, and *Chrm3*, were more frequently expressed in mouse motor cortical neurons than other muscarinic ACh receptors (*Chrm4* and *Chrm5*) and nicotinic ACh receptors (Fig. 6A). In particular, *Chrm3* was expressed in all cell types, whereas *Chrm2* was selectively expressed in the intratelencephalic projection neurons in layer 5 (L5 IT) and two subtypes of GABAergic interneurons, parvalbumin (PV)- and somatostatin (SST)-expressing cells, compared to other subtypes, such as VIP and Lamp5, and other projection neurons in the deep layers (Fig. 6A).

**Figure 6.**
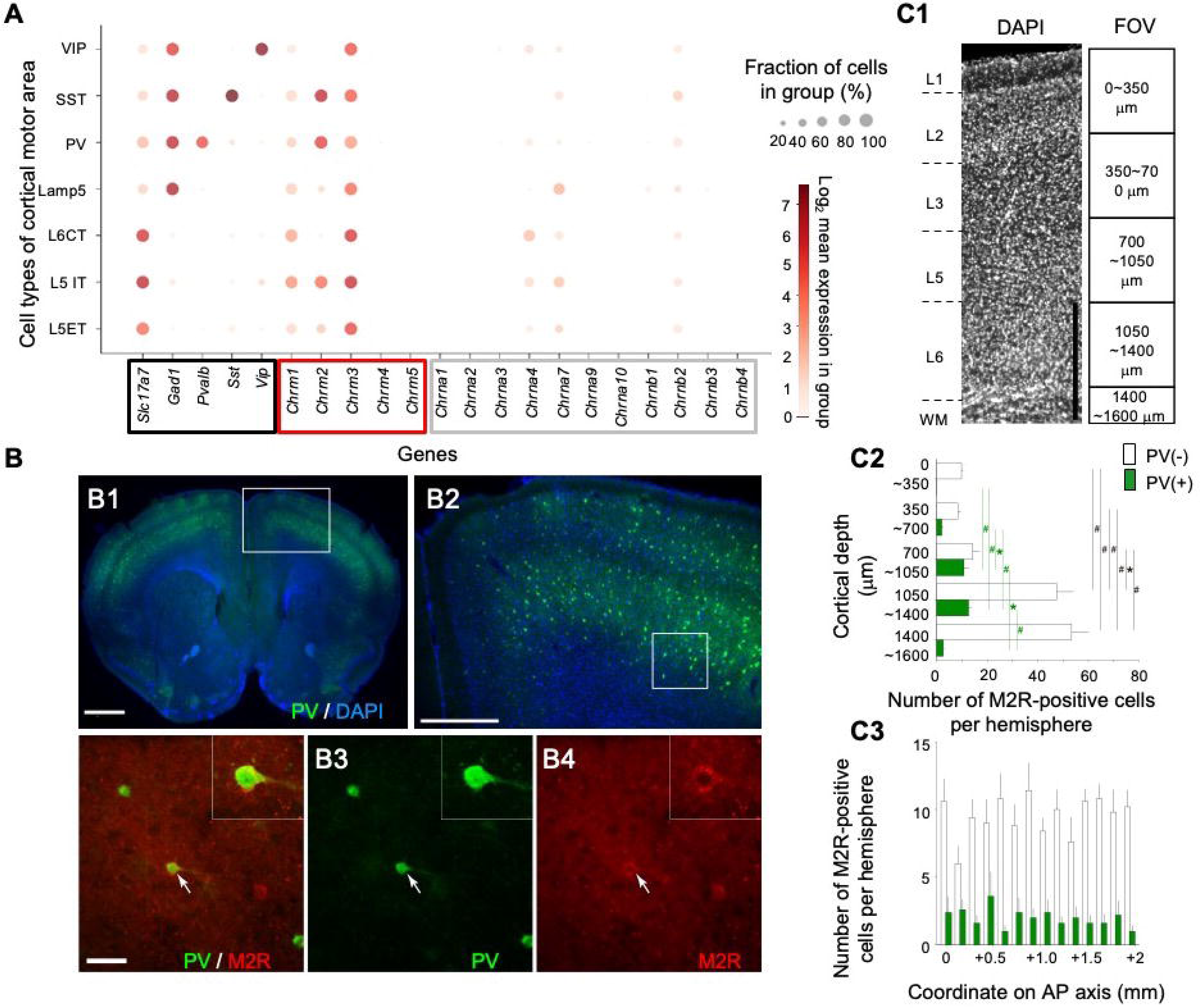
Expression of muscarinic acetylcholine receptor in interneurons of motor cortex. **(A)** Mean expression of mRNA in log scale and fraction of cells expressing mRNA regarding biomarkers of cortical cell-type (black box) and muscarinic (red box) as well as nicotinic (gray box) ACh receptors in cell-types of motor cortical neurons. VIP: vasoactive intestinal peptide, SST: somatostatin, PV: parvalbumin, CT: corticothalamic projection neurons, IT: intratelencephalic projection neurons, ET: extratelencephalic projection neurons. **(B)** B1, A coronal section of the motor cortex showing expression of parvalbumin (PV) and DAPI (blue). Scale bar, 1 mm. B2, A magnified view of the boxed area in B1 showing a distribution of PV-positive cells in layers 2/3 and 5/6. Scale bar, 0.5 mm. B3–B4, Magnified views of the boxed area in B2 showing PV (green) and M2R (red). Representative fluorescence image revealing expression of PV (green) and muscarinic ACh receptor type 2 (M2R) (red). The white arrow indicates a neuron expressing both PV and M2R. Insets, magnified views of a neuron indicated by white arrows. Scale bar, 50 µm. **(C1)** An example of image data analysis. An area in the image of DAPI stained section shows the target position for confocal fluorescent imaging. Field of view (FOV) of the imaging is presented by boxes at the right side of image along cortical depth, but down to white matter (WM). Scalebar, 500 µm. Statistically significant differences among cortical depth were assessed by one-way ANOVA followed by *post hoc* Tukey–Kramer test (*: *P* <0.001, #: *P* <0.0001) for PV-negative (white) and PV-positive (green) M2R-positive cells. (**C2**) The number of PV-negative (white) and PV-positive (green) M2R-posititive cells per each hemisphere along cortical depth. (**C3**) The number of PV-negative and PV-positive M2R-posititive cells per hemisphere along AP axis (ML: 1–2 mm). Values are presented as mean + SEM.

To investigate the expression of the muscarinic ACh receptor type-2 (M2R) encoded by *Chrm2* in GABAergic neurons, we immunolabeled M2R together with PV as a marker of basket cell-like interneurons (Tremblay et al. 2016). We found muscarinic M2R expressing cells in the motor cortex, although M2R-positive cells were sparsely distributed in the motor cortex. We confirmed the presence of double-stained cells in the motor cortex (Fig. 6B). To quantify the number of M2R positive neurons with or without PV expression distributed along the entire cortical depth, we counted M2R positive cells together with checked for the presence or absence of PV across the motor cortex (Fig. 6C1). We found 807 M2R-positive cells in the motor cortex (five hemispheres from three mice), and their distribution was heterogeneous along the cortical depth. M2R-positive cells were more frequently observed at deeper than superficial levels (one-way ANOVA, *P* <10^-7^, *post hoc* Tukey–Kramer test for multiple comparisons). Of the M2R-positive cells, 17.3% were positive cells. Interestingly, M2R/PV double positive interneurons were predominantly distributed in the layers 5 and 6 at a depth of approximately 700–1050 and 1050– 1400 µm, respectively (Fig. 6C2; *post hoc* Tukey–Kramer test for multiple comparison). The ratio of PV-positive M2R cells to PV-negative M2R cells (mean ± SEM, 0.779 ± 0.121) was the largest in the depth ranging between 700–1050 µm for layer 5 (Fig. 6C2). We did not find significant differences in the number of M2R-positive cells along the AP axis (Fig. 6C3, one-way ANOVA, PV-negative M2R cells, *P* = 0.537; PV-positive M2R cells, *P* = 0.863).

These results suggest that ACh could affect more M2R-positive cells, including PV cells, in deep layers than in superficial layers, and M2R-positive cells could be inhibited by ACh because M2R couples to the G_i_ protein; thus, M2R-positive neuronal activity is suppressed by the activation of M2R (Salgado et al. 2007).

## Discussion

The present study revealed that interhemispheric interactions were modulated by reaching behavior and pharmacological cholinergic manipulation. The reaching-related single unit activity revealed much more differential modulation (“reaching-relevance”) between putative cell-types (RS and INT) in contralateral motor cortex than in ipsilateral motor cortex. Reaching-related firing activity in the contralateral motor cortex is significantly altered by paw laterality. The interhemispheric spike correlation was enhanced by the reaching task compared with the non-task period. A negative correlation between contralateral RS and ipsilateral RS neurons and a positive correlation between contralateral RS and ipsilateral FS neurons were enhanced by task performance. The interhemispheric spike correlation between RS and FS neurons was oppositely modulated by the pharmacological upregulation and blockage of ACh. IHI was significantly modulated by the antagonism of muscarinic ACh receptors in deeper cortical layers. The expression of M2R in PV positive cells was more frequently observed in deeper layers than in superficial layers.

### Effect of change in excitatory/inhibitory balance between the homotopic motor cortex on single cell activity

We observed reaching-related neuronal activity in both the contralateral and ipsilateral motor cortical areas, as previously shown in rodent motor cortices during bilateral forelimb pedal tasks (Soma et al. 2017). The relevance was noticeably different between the putative cell types in the contralateral motor cortex. The reaching-related activity of Up-type putative RS cells, assumed to be pyramidal cells (McCormick et al. 1985), and Down-type INT cells, assumed to be GABAergic interneurons (Petersen et al. 2021), was more pronounced in the contralateral motor cortex, potentially contributing to its excitation. Additionally, we identified Up-type and Down-type RS cells in the ipsilateral motor cortex, suggesting these cells can inhibit the homotopic ipsilateral side. This balance of excitatory/inhibitory effects was modulated by paw laterality. In the bilateral paw-reaching task, reaching-related firing activity in contralateral paw reaching was significantly altered in the opposite direction of activity by the opposite paw reaching (*i.e.*, ipsilateral paw reaching), suggesting that the excitatory/inhibitory balance between homotopic motor cortical areas was changed by paw laterality. These results suggest that unilateral paw-reaching activity is modulated by interhemispheric interactions.

### Effect of reaching behavior on interhemispheric spike correlations

Previous physiological studies have shown task- or state-dependent synchronous neural correlations between the bilateral sensory areas (Oran et al. 2021; Adaikkan et al. 2022). Synchronous activity in the interhemispheric homotopic areas is implemented through the corpus callosum (Mohajerani et al. 2010; Oran et al. 2021). We addressed whether reaching behavior also enhanced the neural correlation between homotopic motor cortical areas and found task-dependent enhancement of spike correlation in the STA. During the reaching task, in particular, the STA was negatively enhanced between the contralateral putative RS cells and ipsilateral RS cells, while the STA was positively enhanced between the contralateral putative RS cells and ipsilateral FS neurons. Putative FS neurons are supposed to correspond to the PV-positive GABAergic interneurons (McCormick et al. 1985; Kim et al. 2016). This result suggests that these cellular elements are involved in implementing reaching tasks. Furthermore, the correlation patterns of specific cell pairs could reflect IHI because the conventional IHI circuit model assumes that ipsilateral GABAergic interneurons, which receive transcallosal excitatory inputs, suppress ipsilateral neuronal activity (Toyama et al. 1974; Kawaguchi 1992; Conti and Manzoni 1994; Palmer et al. 2012; Wang et al. 2023).

### Effect of acetylcholine on interhemispheric spike correlations

The change in the interhemispheric spike correlation during task performance may be due to modulation by ACh, which is implicated in regulating arousal and attention (Day et al. 1991; Giovannini et al. 2001; Sarter et al. 2003; Klinkenberg et al. 2011). We found that the interhemispheric synchronous spiking activity of putative RS-FS cell-pairs was differentially altered by pharmacological manipulation of ACh, whereas the ongoing firing rate of both putative RS and FS cells was not. When ACh was upregulated due the inhibition of enzymatic decomposition of ACh, the spike correlation between RS-FS cells was reduced. Conversely, blocking ACh binding to muscarinic ACh receptors resulted in enhanced spike correlation. We hypothesize that callosal pyramidal neurons can be included in the population of putative RS neurons. This result suggests that ACh suppresses the firing of FS neurons only when contralateral putative RS neurons are fired or the temporal correlation between RS and FS neurons becomes less synchronized. In contrast, when ACh was blocked from binding to the muscarinic ACh receptor, the firing activity of putative FS neurons increased only when the putative RS neurons were discharged. Taken together, we suppose that the IHI-related spike correlation could be modulated by ACh.

### Cellular circuitry mechanism underlying cholinergic modulation on IHI

LFP responses to optogenetic excitation of contralateral cortical neurons revealed two components, fast negative and slow positive deflection, reflecting excitatory and inhibitory field potentials, respectively, as previously shown (Spalletti et al. 2017). These fast and slow components were associated with increased and decreased firing rates, respectively, as observed in the mouse auditory cortex (Slater and Isaacson 2020). The fast excitatory and slow inhibitory components of the LFP responses reflect excitatory postsynaptic potentials due to excitatory callosal inputs and feedforward inhibition by GABAergic interneurons (Toyama et al. 1974; Kawaguchi 1992; Conti and Manzoni 1994; Palmer et al. 2012), respectively. We also found that the profiles of the LFP responses to contralateral optogenetic excitation were different at different cortical depths. This transcallosal excitation-inhibition sequence is supported by anatomical evidence. Callosal neurons directly project to contralateral GABAergic interneurons (Karayannis et al. 2007; Rock and Apicella 2015; Anastasiades et al. 2018; Wang et al. 2023). Therefore, the slow inhibitory component may reflect IHI via GABAergic interneurons.

Cholinergic modulation of the slow inhibitory component of the LFP response to contralateral excitation was observed in deep cortical layers. Topical application of a muscarinic cholinergic antagonist enhanced the inhibitory component, suggesting that contralateral GABAergic cells could be excited in response to callosal input in deep layers 5 and 6 and that, in normal situations, ACh could exert an inhibitory action on contralateral GABAergic interneurons through muscarinic ACh receptors. PV neurons in the deep layers of the frontal cortex receive direct callosal innervation from the contralateral hemisphere (Karayannis et al. 2007). Furthermore, a subtype of the muscarinic ACh receptor, M2R, was expressed in deep layers, and approximately 17% of M2R-positive neurons expressed PV. Previous studies have suggested that GABAergic neurons expressing cholinergic receptors and receiving cholinergic inputs modulate cortical activity depending on their behavior (Fu et al. 2014; Kuchibhotla et al. 2017; Ren et al. 2022). This result supports our physiological data that the muscarinic receptor antagonist enhances the inhibitory component of the LFP response to contralateral excitation in deep layers because M2R couples to the G_i_ protein, and thus could suppress M2R-positive neurons (Gu 2002; Salgado et al. 2007). Taken together, these results suggest an IHI circuitry model that is modulated by ACh. The IHI neural circuit consists of GABAergic neurons in deep layers that receive direct callosal excitatory inputs from the contralateral homotopic motor cortex, and some of the GABAergic neurons expressing muscarinic ACh receptors, which are coupled to the G_i_ protein. As ACh release occurs, GABAergic neurons are inhibited via muscarinic receptors, resulting in the disinhibition (excitation) of postsynaptic neurons.

## Acknowledgements

This work was supported by JSPS KAKENHI 19H0572301, 21H02581 and 23K18256 to HA, KAKENHI 22K06485 to TH, and the Program of the Network-type Joint Usage/Research Center for Radiation Disaster Medical Science. We thank Prof. Atsu Aiba for Emx1-Cre mouse line, Y. Ishimaru, F. Nishimura, S. Wakida, and the Natural Science Center for Basic Research and Development of Hiroshima University for their technical support.

## Conflict of interest

The authors declare no competing financial interests.

## Notes

### Competing Interest Statement

The authors have declared no competing interest.

